# LEN-Seek: Fast and scalable ligand binding-site similarity search in the latent space of an SE(3)-invariant graph VAE

**DOI:** 10.64898/2026.08.14.744759

**Authors:** Kyunghwan Yeo, Dongwoo Kim, Jaemin Sim, Juyong Lee

## Abstract

**Motivation:** Ligand binding-site similarity search is a crucial step in drug discovery that reduces the conformational search space for docking and other downstream tasks by comparing a target protein against experimentally identified binding sites. Existing methods rely on either direct structural alignment or lossy compression of structural information, producing a trade-off between scalability and precision.

**Results:** We propose LEN-Seek, a ligand binding-site search method based on a graph neural network (GNN)-driven variational autoencoder (VAE) that encodes the 3D structural and physicochemical context of a binding site into a probabilistic latent space, enabling similarity search within a low-dimensional vector space. A binding site is modeled as a graph of amino acid residues, with node features adopted from the protein language model, Ankh, and edges encoded as SE(3)-invariant (roto-translational invariant) geometric relationships, thereby avoiding expensive data augmentation or SE(3)-equivariant models. Compared to ProBiS, the purely geometric graph-clique based method, LEN-Seek successfully retrieves a substantial portion of similar binding sites with a roughly 3,400-fold lower per-comparison cost, demonstrating its potential as a scalable approach to template-based ligand binding-site search in large-scale protein structure databases.

**Supplementary information:** Supplementary data are available at Bioinformatics online.

## Introduction

Discovery of novel drug candidates is a pivotal yet time-consuming and capital-intensive challenge in modern medicine[12]. Computational drug discovery screens hit candidates in advance through virtual simulation, improving the efficiency of the development process[7, 53]. Molecular docking is a core step that identifies candidate ligands and predicts their bound conformation and affinity. Blind docking over the entire protein surface is computationally expensive and prone to false positives, whereas targeted docking with a predefined binding site markedly improves both efficiency and accuracy[9]. Co-folding methods such as AlphaFold3[1], Boltz-1[59], and RoseTTAFold All-Atom[41] likewise benefit from prior binding-site information. Accurate identification of a target’s ligand binding site is therefore a prerequisite for successful computational drug discovery and for structure-based drug design[2, 4].

### Binding Site Identification and the Scalability–Precision Trade-off

Binding site prediction methods are generally categorized as geometry-, energy-, or template-based[39, 64]. Geometry-based methods locate cavities from surface shape, such as SURFNET[43], LIGSITE[21, 24], FPocket[45], and have evolved into learning-based predictors that learn the geometric patterns of surface points, voxels, atoms, or residues, including P2Rank[42], Kalasanty[55], PUResNet[29], SwinSite[31], PointSite[60], EquiPocket[63]. Energy-based methods place chemical probes on the surface and seek free-energy minima (Q-SiteFinder[44], FTMap[38]), offering physical validity at high computational cost. Template-based methods instead search a database of experimentally verified binding sites for similar ones, inheriting the biological validity of those experimental sites[64, 39]. These approaches rest on the hypothesis that binding sites with similar structure and physicochemical properties bind similar ligands, supported by molecular complementarity[49, 37], convergent evolution of catalytic geometries[40], and drug-promiscuity statistics—e.g. about 71% of multi-target drugs bind structurally similar sites while 55% lack global structural similarity[23, 20].

Traditionally, graph-based alignment tools quantify binding-site similarity accurately: ProBiS[36, 35] aligns surface graphs via maximum-clique search[8], APoc[18] introduces the PS-score, and G-LoSA[46] compares chemical feature points with a size-normalized GA-score. However, their one-by-one alignment scales poorly: the PDB now exceeds 240,000 structures[6] and the AlphaFold Database over 200 million[57], making pairwise alignment intractable at database scale. Faster alternatives simplify the representation. Geometric hashing (SiteEngine[51]) and 1D distance distributions (PocketMatch[61]) lose spatial detail, and Foldseek[56] encodes backbone geometry as a 1D sequence but is ill-suited to binding sites, which are discontinuous residue sets. Machine-learning methods learn binding-site representations directly: DeeplyTough[52] encodes voxel grids with a 3D CNN and MaSIF[17] encodes surface meshes, but both lose information during discretization and require costly rotation augmentation. More fundamentally, 3D structures are roto-translationally equivariant; equivariant networks such as SE(3)-Transformer[16] and EGNN[50] avoid augmentation but rely on expensive operations that hinder large-scale parallel computation. A method is therefore needed that searches similar binding sites at high speed regardless of global fold, minimizing the scalability–precision trade-off.

### Technical Goals

Here, we propose LEN-Seek (Latent Encoding Network for template Seeking), a method based on a graph neural network (GNN) and a variational autoencoder (VAE)[32] that compresses the structural and physicochemical information of a binding site into a low-dimensional, continuous probabilistic latent space, replacing partial-structure alignment with distance computation in latent space. Our method pursues three goals. First, a continuous latent space: unlike discrete or ordinary autoencoder representations, a VAE places encoded data on a continuous manifold so that proximity in latent space reflects genuine structural and physicochemical similarity[32]. Second, physicochemical encoding via representation learning[5]: instead of hand-crafted property rules, we use the protein language model (PLM) Ankh[48, 14] to capture evolutionary and physicochemical context directly from sequence. Third, geometric robustness and efficiency through SE(3)-invariant (roto-translational invariant) features: following Ingraham *et al*.[25]—and unlike RoseTTAFold2[3], which retains an SE(3)-equivariant module, or SimpleFold[58], which relies on random rotation augmentation— we represent 3D structure with local-frame-based invariant relations from the input stage, avoiding equivariant networks and data augmentation. Together these aim to deliver scalable, structure- and chemistry-aware binding-site search for large-scale databases.

## Methods

### Overview

LEN-Seek is a deep learning approach built on a VAE, as illustrated in Figure 1. The model is trained to take ligand binding-site data as input and reconstruct the original binding site. Once trained, the encoder extracts latent vectors from the ligand binding sites in the training data to construct a latent-space dataset. When a novel ligand binding site is given as a query, the encoder transforms it into a set of latent vectors, and a similarity search is then performed within the latent-space dataset to retrieve similar ligand binding sites. Each binding site enters the model as a graph whose node features are built from Ankh (Figure 1B).

**Figure 1.**
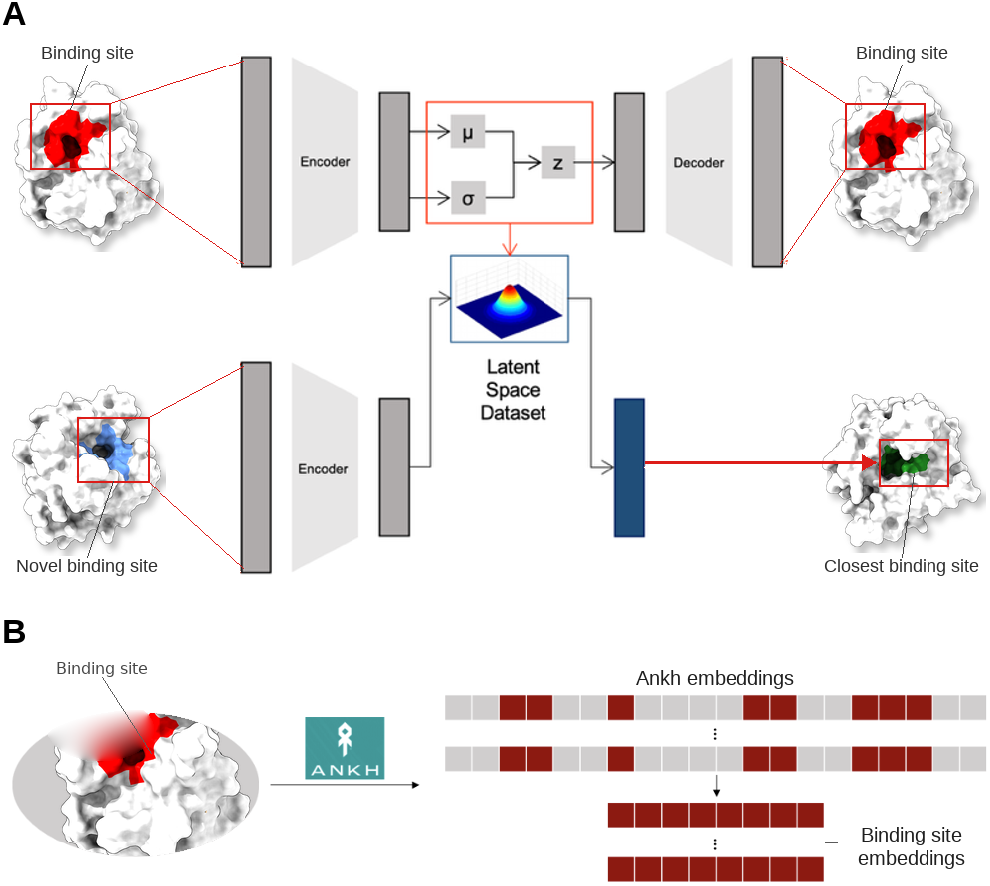
Overview of LEN-Seek. A. The graph VAE is trained to encode and reconstruct ligand binding sites. Once trained, the encoder maps the training binding sites into a latent-space dataset that serves as the search space; a novel query binding site is encoded in the same way and compared against this dataset to retrieve the most similar sites. B. Construction of the node features that feed the encoder in A. The full protein sequence is passed through the protein language model Ankh to obtain a 768-dimensional per-residue embedding, and the embeddings of the residues constituting the binding site are then collected to form the node features (the binding-site embeddings).

The encoder maps each residue of a binding site into a probabilistic latent vector, turning the site into a set of latent vectors lying on a continuous distribution, while the decoder reconstructs the input from them, inducing latent representations that reflect structural and physicochemical context. At evaluation, only the encoder is used: training data are encoded into a latent-space dataset that serves as the search space, and a query site is encoded into its residue-level latent set for distance-based retrieval.

### Database

We used our in-house BsitePDB, a calibrated set of ligand binding sites from the PDB[30]. BsitePDB consists of 163,252 ligand binding sites from 66,912 proteins, drawn from PDB entries deposited up to November 6, 2023. Only binding sites meeting the definition of a ligand binding site were included, following criteria from sc-PDB[11] and ProBiS-Database[34]. The criteria for defining a ligand binding site are outlined in Supplementary Table S1.

Data registered in the PDB after July 21, 2023 were held out as the evaluation dataset. During redundancy filtering with MMseqs2[54], entries sharing over 80% sequence similarity with any training data were excluded, resulting in a final set of 482 ligand binding sites from 161 proteins.

### Roto-Translational Invariant Representation

A ligand binding site is interpreted as a set of amino acid residues, which is represented as a geometric graph. In a graph, each residue is a node whose feature is a PLM embedding capturing its evolutionary and physicochemical context. Each edge encodes the SE(3)-invariant relative geometry between two residues, so that residue chemistry is carried by the nodes and binding-site structure by the edges. To address the roto-translational equivariance inherent in 3D structures, the entire graph is constructed using only roto-translational invariant features, expressing the structure as relative geometric relationships between residues. This eliminates the need for computationally heavy operations such as data augmentation or equivariant neural networks. The features of the binding-site graph are computed from three basic elements: the PLM Ankh embedding (**h**), the local frame of each amino acid (**R**), and the Cartesian coordinate of its alpha carbon 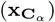.

Here **h**_*i*_ ∈ ℝ^768^ is the Ankh embedding, 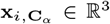 the alpha-carbon coordinate, and **R**_*i*_ ∈ SO(3)[19] the local frame, where SO(3) is the group of rotations about the origin in 3D space. The number of graph nodes was fixed at 65 for all data (*i* ∈ {1, …, 65}). As shown in Supplementary Figure S1, the largest binding site in BsitePDB contains 65 residues, which motivated standardizing the node count to this value for efficient data processing. For each data point, all values associated with padded nodes were set to 0 and masked so that they did not participate in any computation. Sites with fewer than 10 nodes were excluded, as too few residues make it difficult to capture geometric context, leaving a total of 114,662 data points for training.

#### Edge Feature: Structure Representation

The relative geometric relationships between the amino acids constituting a ligand binding site are encoded as edge features in the graph. The edge feature connecting two nodes *i* and *j* is a 39-dimensional vector **e**_**ij**_ that combines three elements, as shown in eq. (1): a 32-dimensional Radial Basis Function (RBF) kernel embedding *ϕ*_dist_ (**d**_*ij*_) of the Euclidean distance **d**_*ij*_ between the two residues, a 3D unit direction vector **o**_**ij**_, and a 4D quaternion rotation vector **q**_**ij**_ between the two local frames.

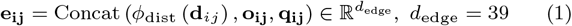

Because absolute atomic coordinates change under global rotation and translation, we attach a local coordinate frame to every residue and express all inter-residue geometry relative to these frames; any quantity measured this way is therefore invariant to the binding site’s overall position and orientation. Concretely, a local frame **R**_**i**_ was defined from the backbone coordinates of the amide nitrogen (**x**_**i**,**N**_), alpha carbon 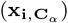, and carbonyl carbon (**x**_**i**,*C*_) of each residue. The first axis **u**_1_ is the unit vector from **C**_*α*_ to **C**; the second axis **u**_2_ is the Gram–Schmidt-orthogonalized direction toward **N**; and the third is **u**_3_ = **u**_1_ × **u**_2_, giving the local frame **R**_**i**_ = [**u**_1_, **u**_2_, **u**_3_] ∈ SO(3). The distance between residues is computed from the alpha-carbon coordinates, 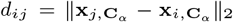. This distance is an inherently roto-translational invariant scalar that requires no additional treatment, but it was projected into a 32-dimensional vector space via an RBF kernel so that the model can flexibly learn weights as a function of distance. Each component of this 32-dimensional vector represents the probability that the actual distance falls in a normal distribution, defined by dividing the range 0–20 Å into 32 intervals, with each interval’s median as the mean and its width as the standard deviation, yielding *ϕ*_dist_(*d*_*ij*_) ∈ ℝ^32^. The direction between residues is likewise computed from the alpha-carbon coordinates. Although a direction vector in 3D Euclidean space is roto-translational equivariant, projecting it onto each residue’s local frame, as shown in eq. (2), makes it roto-translational invariant. A small constant *ϵ* was added for numerical stability.

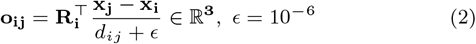

Relative rotation between two local coordinate systems is expressed as a 4-dimensional quaternion **q**_**ij**_ = Quat(**R**_**i**_^⊤^**R**_**j**_) ∈ ℝ^4^.

The final 39-dimensional edge feature vector was formed by concatenating the 32-dimensional distance vector, the 3-dimensional direction vector, and the 4-dimensional rotation vector obtained above. Edges were defined only between the 15 nearest nodes, so that the model learns only local relationships within the ligand binding site.

#### Node Feature: Protein Language Model

Each amino acid is represented as a single node, and only Ankh PLM embeddings were used as node features. The Ankh embeddings were computed after separating all chains of the protein. As shown in Figure 1B, only the embedding vectors of the ligand binding site residues were parsed from the full set of embeddings.

### Model Architecture

The LEN-Seek model follows a Graph Transformer-based VAE structure[13], as shown in Figure 2. It is designed to simultaneously process the geometric structure and high-dimensional physicochemical features of a binding site, encoding them into a low-dimensional, continuous latent space and then reconstructing the original structure and features.

**Figure 2.**
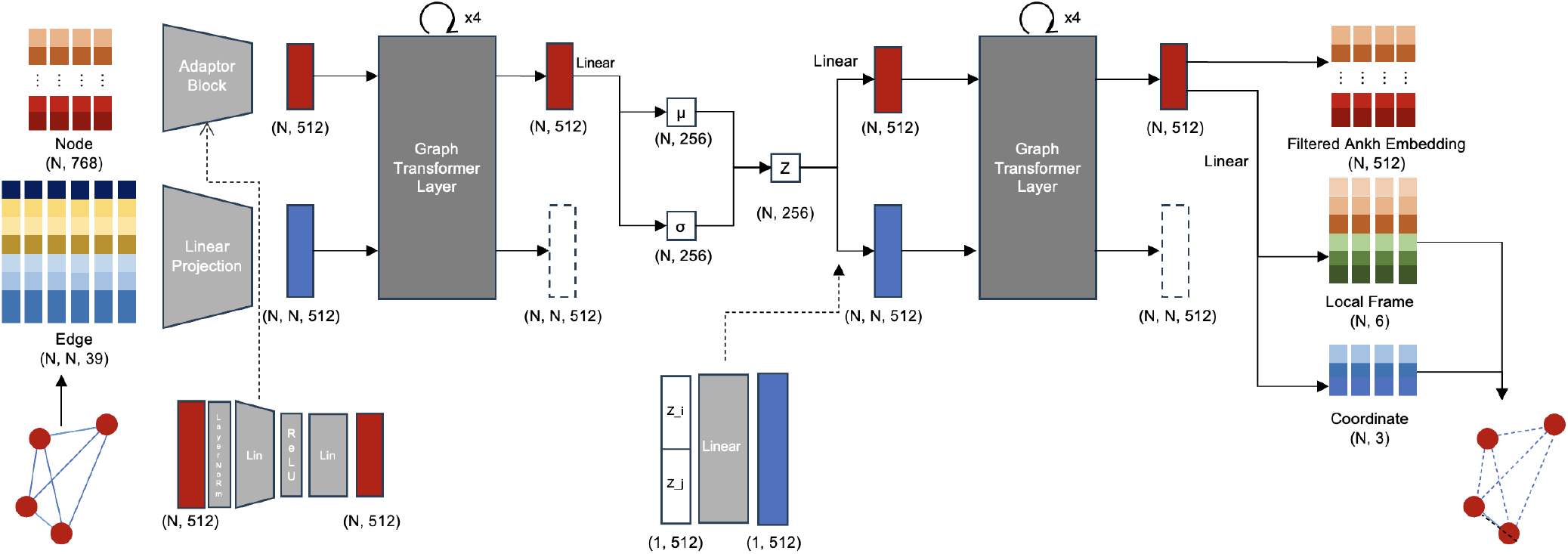
Model architecture of LEN-Seek. The model is based on a graph transformer. After passing through the Adaptor Block, the node and edge features are mutually updated across four Graph Transformer layers. Only the updated node features are used to map the data onto the latent space, while the edge data are discarded to prevent leakage of connectivity information to the decoder. The reparametrized node features are then used to generate new edge data, which pass through four Graph Transformer layers to reconstruct the original ligand binding site.

The edge and node features of the input first pass through an ‘Adaptor Block’. The 768-dimensional Ankh embedding is projected into a 512-dimensional space by an MLP with ReLU activation, which ‘adapts’ the high-dimensional feature space of the PLM so that the model can better learn the structural context of the ligand binding site. The edge features are simultaneously projected to 512-dimensional initial edge representations through a single linear layer.

The encoder consists of four graph attention layers. Unlike standard Transformers that consider only node–node relationships, we integrate the edge features directly as a bias in the attention computation[62]. This approach, also used in geometric deep learning models such as the Evoformer module[27] of AlphaFold, imposes geometric constraints that directly regulate the strength of information propagation.

In the *l*-th update, per-head queries are derived from node *i* and keys and values from node *j* (each a linear map of the LayerNorm-ed node feature, head dimension *d*_*k*_ = 64). The key novelty is that edge information enters as a bias term *b*_*ijk*_ added to the attention logits, as shown in eqs. (3)–(4).

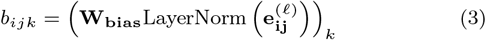

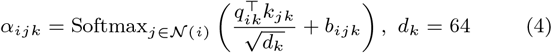

where **W**_**bias**_ is a learned projection of the LayerNorm-ed edge feature 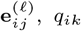 and *k*_*jk*_ are the query of node *i* and the key of node *j* for head *k, α*_*ijk*_ is the resulting attention weight, and *N* (*i*) is the neighborhood of node *i*.

Node information is updated by summing the values of the connected nodes weighted by the attention values, and the updated node information 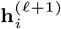 is then obtained through a residual connection over *N*_head_ = 8 attention heads. Edge information is then updated from the LayerNorm-ed edge feature concatenated with the two updated node features through an MLP with a residual connection.

Among the final encoder outputs, only the updated node information 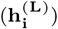 is mapped, through linear layers, to a 256-dimensional mean (*µ*_**i**_) and log-variance 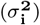 vector for each node, as in eq. (5); the edge information 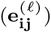 is discarded to prevent direct leakage of connectivity to the decoder. The input graph is thereby placed on a latent space, and the latent vector (**z**_**i**_) is generated by adding noise *ϵ* drawn from a standard normal distribution via the reparameterization trick[32], which becomes the input to the decoder.

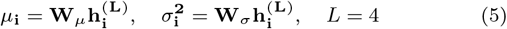

The decoder is trained to reconstruct the structure and features of the original ligand binding site from node information 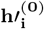 and edge information 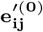 created from the latent vectors. Like the encoder, the decoder consists of four graph attention blocks, but its input edge information is newly created by concatenating the latent vectors of connected nodes and passing them through a linear layer. The decoder operates on a fully connected graph.

The output layer of the decoder branches into three heads that recover the node feature 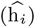, the alpha-carbon coordinate 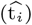, and the local-frame axes 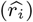 of each residue. For the node feature, the model recovers the 512-dimensional adapted vector rather than the entire original Ankh embedding. For the local frame, since the third axis is fully determined by the other two, only two axes are reconstructed. The three decoder heads output 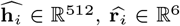, and 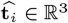, respectively.

### Loss Function

The model’s loss combines a reconstruction loss (L_recon_) and a KL-divergence term (L_KL_), following the conventional VAE objective, as shown in eq. (6).

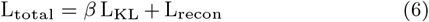

The variational posterior predicted by the model is *q* (**z**_**i**_|*G*), the conditional distribution given the graph *G*. The KL term drives this posterior toward the prior *p* (**z**_**i**_), the distribution that the node-wise information constituting the ligand binding site is assumed to follow. When minimizing the KL term, a VAE can collapse the posterior toward the prior and behave like an ordinary autoencoder—the so-called “posterior collapse” phenomenon. To prevent this, the KL weight was set to 0 at the start of training so the model could first focus on reconstruction. In addition, the Free Bits strategy[33] was applied to encourage better learning of the distribution; it enforces a lower bound so that the KL loss cannot fall below a set threshold, ensuring the latent variables retain a minimum amount of information. This threshold was set to 12.

The reconstruction loss consists of a node-feature reconstruction loss (L_feat_) and a structure reconstruction loss (L_struct_), as defined in eq. (7), to ensure recovery of both physicochemical and structural information.

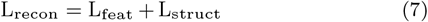

The node-feature reconstruction loss L_feat_ combines three terms: an MSE term (L_MSE_) for numerical recovery, a cosine-similarity term (L_Cos_) for direction recovery, and an L2-norm term (L_Norm_) for scale recovery. To compensate for their differing value ranges, weights were applied, as shown in Supplementary Table S2.

For the structure reconstruction loss, we adopted the FAPE (Frame-Aligned Point Error) loss (L_FAPE_) introduced in AlphaFold[27]. It measures the relative positional error of the other residues in each residue’s local frame, making it roto-translational invariant. An RMSD (Root-Mean-Square Deviation) loss (L_RMSD_) was additionally applied to the Cartesian coordinates predicted by the model. Because coordinates are roto-translational equivariant, the rotational and translational components were removed by Kabsch alignment[28] before computing the RMSD. The structure loss L_struct_ is a weighted sum of the FAPE and RMSD terms. Because the loss terms span different value ranges, their weights were tuned accordingly; the weights and hyperparameters used for training are detailed in Supplementary Table S2 and Supplementary Figure S2.

## Results

### Feature and Coordinate Reconstruction

To determine whether trained model successfully learned the structure and Ankh embedding context, we first verified the model’s ability to recover original data. BsitePDB evaluation dataset was utilized for reconstruction performance assessment.

For the Ankh embeddings, recovery was evaluated on the 512-dimensional vectors processed through the adaptation module. The model achieved successful recovery, with an average cosine similarity of 0.994 (Figure 3A). A Principal Component Analysis (PCA) of the adapted 512-dimensional embeddings confirmed that the distribution was preserved after reconstruction (Figure 3B).

**Figure 3.**
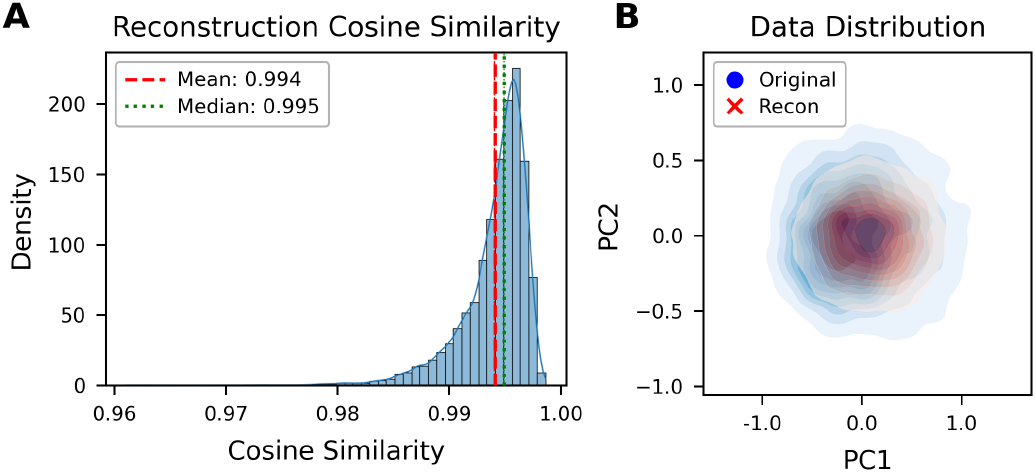
Reconstruction of node feature. A. Cosine similarity between the original ‘adapted’ Ankh embedding and the reconstructed vector. B. Distribution of the first two principal components of the ‘Adapted’ 512-dimensional Ankh embedding. The original distribution is shown in blue and the reconstructed distribution in red.

For structure, the model successfully reconstructed the binding-site geometry, with an average FAPE of 2.10 Å and RMSD of 1.07 Å (Supplementary Figure S3).

### Similarity Search

To demonstrate that LEN-Seek identifies similar ligand binding sites in latent space, large-scale similarity-search experiments were conducted on the BsitePDB evaluation data. LEN-Seek performed the searches after constructing a ‘Latent Space Database’ by mapping the training data into latent space with the encoder.

Since a ligand binding site is a variable-size set of latent vectors, one per amino acid residue, standard metrics such as Euclidean distance or cosine similarity are unsuitable. Treating each residue’s latent vector as a point allows a binding site to be represented as a point cloud. We therefore used the L2 Chamfer distance[15] as the similarity metric, as it measures the dissimilarity between two sets regardless of their sizes, as defined in eq. (8).

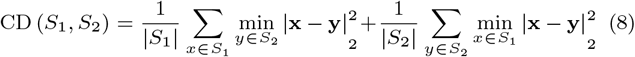

While existing structure alignment tools like ProBiS ensure high accuracy, one-by-one comparison across all protein databases has clear scalability limits. Thus, LEN-Seek was utilized as a primary high-speed filtering tool to select candidate groups, which were then cross-validated using ProBiS. We adopted a two-tier evaluation. ProBiS and G-LoSA were treated as established structure-alignment baseline methods—deterministic algorithms rather than learned models, but still only one structural criterion each and not absolute ground truth—against which we measured concordance, whereas ligand chemical similarity (see the independent validation below) provides an alignment-free criterion that directly tests the biological hypothesis underlying template search. Consequently, the confusion-matrix statistics reported here (true/false positives and negatives, precision, recall, F1, and the Matthews correlation coefficient, MCC) are defined *relative to the ProBiS baseline* and quantify agreement with it rather than absolute correctness. Following this convention, a ProBiS Z-score of 2.0 or higher was defined as a significant alignment (‘Positive’), and lower scores as non-significant (‘Negative’)[34].

Searches were performed using the Chamfer distances computed by LEN-Seek for the 482 binding sites in the BsitePDB evaluation data. The optimal similarity threshold was determined from the MCC[10] against the ProBiS results. A Chamfer distance of 50.0, which yields the highest MCC for the top-1 results, was selected (Supplementary Figure S4).

The top-ranked results and statistics are presented in Table 1. LEN-Seek achieved precision and recall above 0.80 through the top-50 results. Even on this unbalanced data, where dissimilar sites outnumber similar ones, it maintained an F1-score above 0.8 through the top-50 and an MCC above 0.6 through the top-100, demonstrating strong concordance with the ProBiS.

**Table 1.** Concordance of LEN-Seek top-ranked search results with the ProBiS baseline. Cases with a ProBiS Z-score over 2.0 (the threshold for significant alignment) were treated as similar (‘Positive’), others as ‘Negative’; for LEN-Seek a Chamfer distance below 50.0 was treated as similar. True/false positives and negatives are defined relative to this ProBiS baseline, not as absolute ground truth. Counts can slightly exceed 482 × *k* because binding sites with tied Chamfer distances share a rank and are all retained.

| Rank | TP | FP | FN | TN | Precision | Recall | F1-Score | MCC |
| --- | --- | --- | --- | --- | --- | --- | --- | --- |
| 1 | 306 | 31 | 25 | 120 | 0.9080 | 0.9245 | 0.9162 | 0.7273 |
| 5 | 1378 | 148 | 190 | 693 | 0.9030 | 0.8788 | 0.8908 | 0.6953 |
| 10 | 2525 | 327 | 435 | 1538 | 0.8853 | 0.8530 | 0.8689 | 0.6713 |
| 20 | 4462 | 712 | 959 | 3511 | 0.8624 | 0.8231 | 0.8423 | 0.6512 |
| 50 | 8996 | 2194 | 2090 | 10819 | 0.8039 | 0.8115 | 0.8077 | 0.6425 |
| 100 | 13986 | 5235 | 3752 | 25228 | 0.7276 | 0.7885 | 0.7568 | 0.6073 |

For the top-100 results, the ProBiS Z-score and the LEN-Seek Chamfer distance showed a Pearson correlation of −0.464 and a Spearman correlation of −0.419 (Figure 4A). Because the Z-score has limitations (it is a statistical significance indicator and is undefined for non-significant alignments), we additionally analyzed the GA-score from G-LoSA across all comparisons; this yielded a Pearson correlation of −0.853 and a Spearman correlation of −0.826 (Figure 4B), indicating a strong association with existing geometric algorithm-based structure-alignment methods.

**Figure 4.**
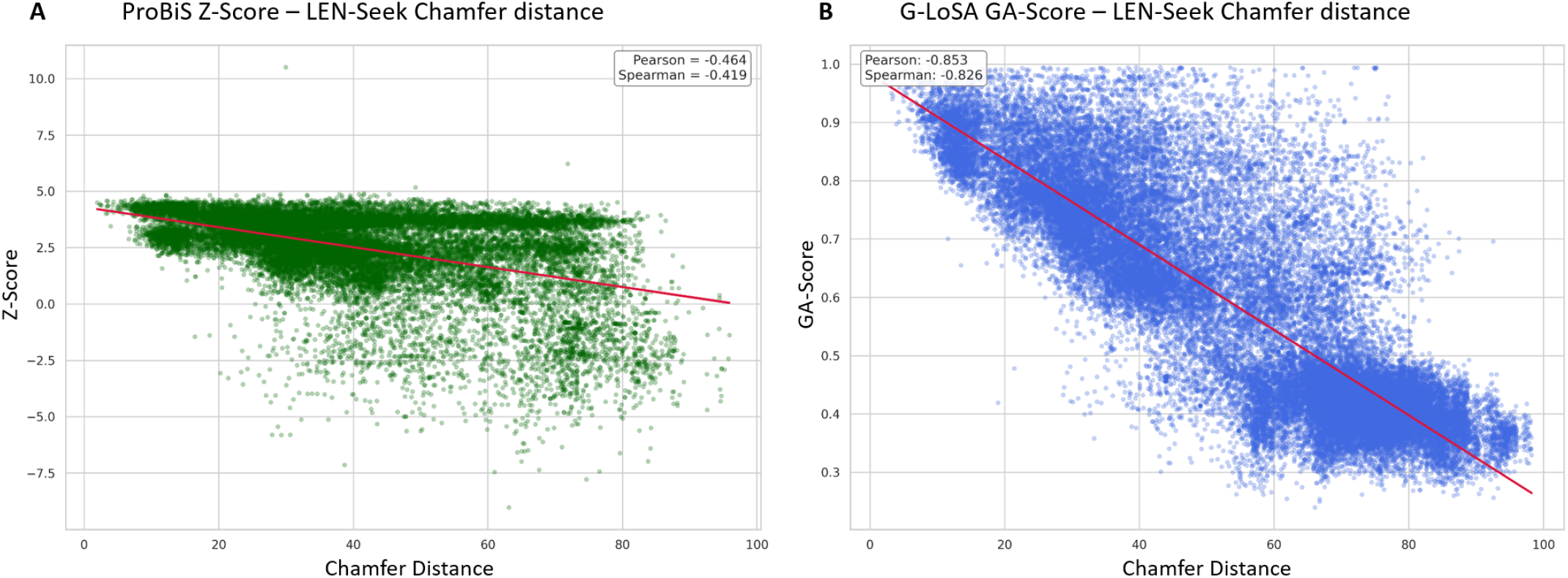
Correlation between LEN-Seek’s Chamfer distance and the ProBiS Z-score / G-LoSA GA-score for LEN-Seek’s top-100 search results. A. ProBiS Z-score (vertical axis) versus LEN-Seek’s Chamfer distance (horizontal axis). Note that the Z-score expresses the statistical significance of similarity between binding sites rather than a quantitative measure of similarity. B. G-LoSA GA-score (vertical axis) versus LEN-Seek’s Chamfer distance (horizontal axis).

### Independent Validation by Ligand Similarity

Concordance with ProBiS (above) measures agreement with one structure-alignment method, not biological correctness. We therefore validate LEN-Seek against an *independent*, alignment-free criterion: the chemical similarity of the ligands that the retrieved sites actually bind. This directly tests the core hypothesis of template-based search—that “binding sites with similar structures and physicochemical properties bind similar ligands”—through a criterion that is independent of structural alignment. Ligand similarity comparisons were conducted on pairs classified as similar or dissimilar by LEN-Seek and ProBiS, using experimentally validated binding ligands from PDB.

Ligand similarity was measured as the Dice similarity of 2048-bit Morgan fingerprints[47], which encode bonded connectivity and reflect the topological environment and substructure overlap. Using the classification criteria of Chamfer distance 50.0 for LEN-Seek and Z-score 2.0 for ProBiS, the ligand-similarity distributions were approximated by kernel density estimation (Figure 5). A Tanimoto similarity of 0.3 is an established indicator of significant activity association[26], which corresponds to a Dice similarity of approximately 0.46 (*D* = 2*T/*(1 + *T*)).

**Figure 5.**
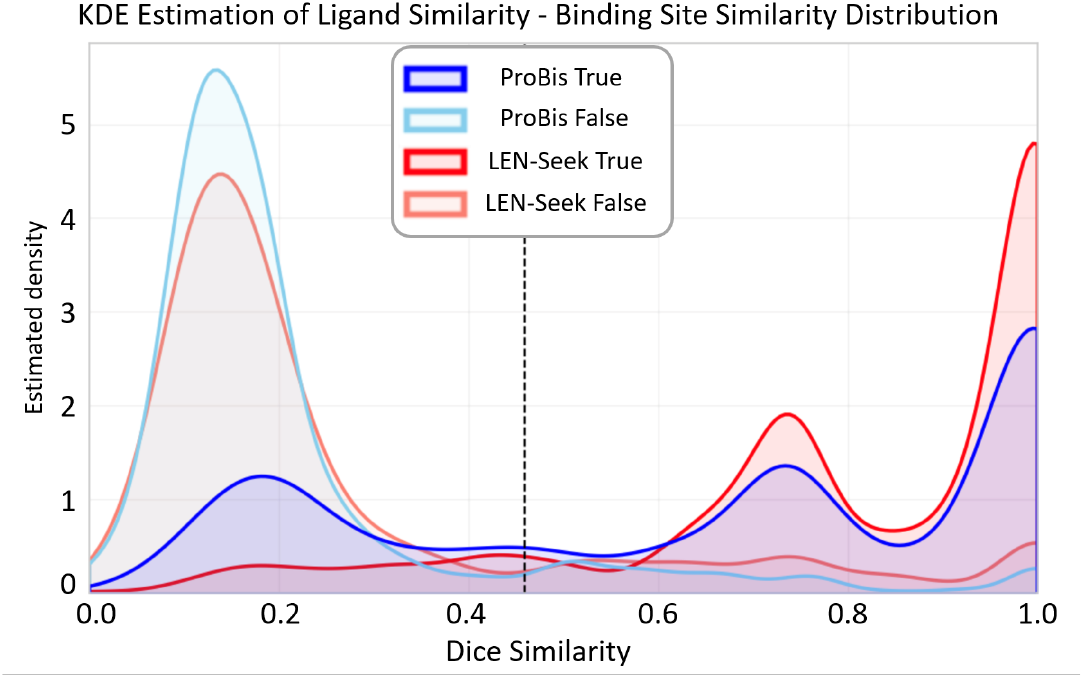
Kernel density estimates of the ligand-similarity distributions for the LEN-Seek and ProBiS results.

Among the pairs exceeding this threshold, 88.0% of LEN-Seek’s similar pairs qualified, versus 19.09% of its dissimilar pairs, a clear separation; for ProBiS the corresponding values were 68.45% and 10.76%. The proportion of similar ligands among the identified pairs was thus higher for LEN-Seek than for ProBiS. At a stricter Dice similarity of 0.6 (Supplementary Table S3), LEN-Seek likewise showed a large gap between similar (84.64%) and dissimilar (14.49%) pairs, confirming agreement with the core hypothesis of structure-based search.

### Computational Efficiency

Screening times for ProBiS, G-LoSA, and LEN-Seek were measured on 92 ligand binding sites from 50 proteins in the evaluation set. ProBiS and G-LoSA were run against all 163,252 training sites 16 Intel Xeon Gold 6348 CPUs, giving 92 × 163,252 = 15,019,184 comparisons, whereas LEN-Seek was run against the 114,662 sites retained during training-data generation on a single Nvidia GeForce RTX 4090 GPU, giving 92 × 114,662 = 10,548,904 comparisons.

Comparison results are detailed in Table 2. ProBiS and G-LoSA took approximately 0.443 and 0.622 seconds per comparison on 16 CPU cores, respectively, whereas LEN-Seek averaged 0.00013 seconds per comparison on a single GPU (1387 seconds for 10,548,904 comparisons). We emphasize that this roughly 3,400-fold per-comparison gap is not hardware-normalized: the methods were run on their respective recommended hardware (16 CPUs vs. one GPU) and over slightly different database sizes (163,252 vs. 114,662 sites). LEN-Seek additionally incurs a one-time preprocessing cost (Table 2; ≈13,000 s in total, dominated by training-set Ankh embedding). This cost is amortized across queries and is small relative to alignment-based search: ProBiS scans the full database in ≈0.443s × 163,252≈7.2×10^4^ s per query, so even when preprocessing is included LEN-Seek is faster from the first query, and the per-query advantage grows with the number of queries.

**Table 2.**
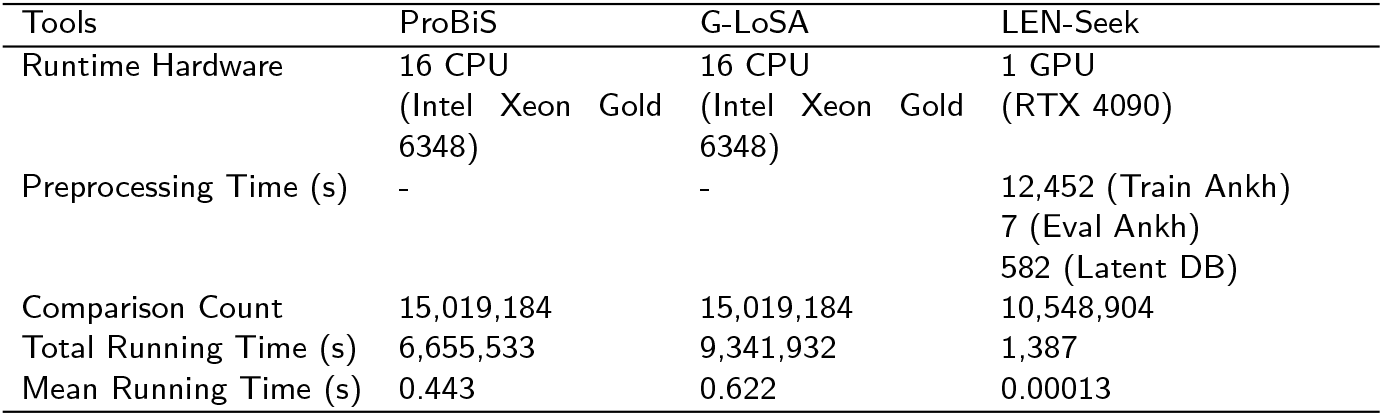
A comparison of computational time of ProBiS, G-LoSA, and LEN-Seek.

| Tools | ProBiS | G-LoSA | LEN-Seek |
| --- | --- | --- | --- |
| Runtime Hardware | 16 CPU<br>(Intel Xeon Gold 6348) | 16 CPU<br>(Intel Xeon Gold 6348) | 1 GPU<br>(RTX 4090) |
| Preprocessing Time (s) | - | - | 12,452 (Train Ankh)<br>7 (Eval Ankh)<br>582 (Latent DB) |
| Comparison Count | 15,019,184 | 15,019,184 | 10,548,904 |
| Total Running Time (s) | 6,655,533 | 9,341,932 | 1,387 |
| Mean Running Time (s) | 0.443 | 0.622 | 0.00013 |

## Discussion

LEN-Seek achieved strong retrieval among top-ranked results in large-scale similarity searches over massive protein databases; in some cases, however, its results diverged from those of the structure-alignment-based ProBiS. By analyzing these discrepancies, we sought to understand how this deep-learning approach perceives binding sites and how it differs from conventional, purely geometric methods.

### Shape Sensitivity

Cases classified as ‘False Negatives’—where LEN-Seek indicated low similarity (large Chamfer distance) while ProBiS indicated high similarity (high Z-score)—were analyzed across three scenarios: identical binding ligands, similar ligands, and dissimilar ligands (Figure 6). Ligand similarity was determined using 2048-bit Morgan fingerprint vectors [47], considering Dice similarity over 0.7 as similar.

**Figure 6.**
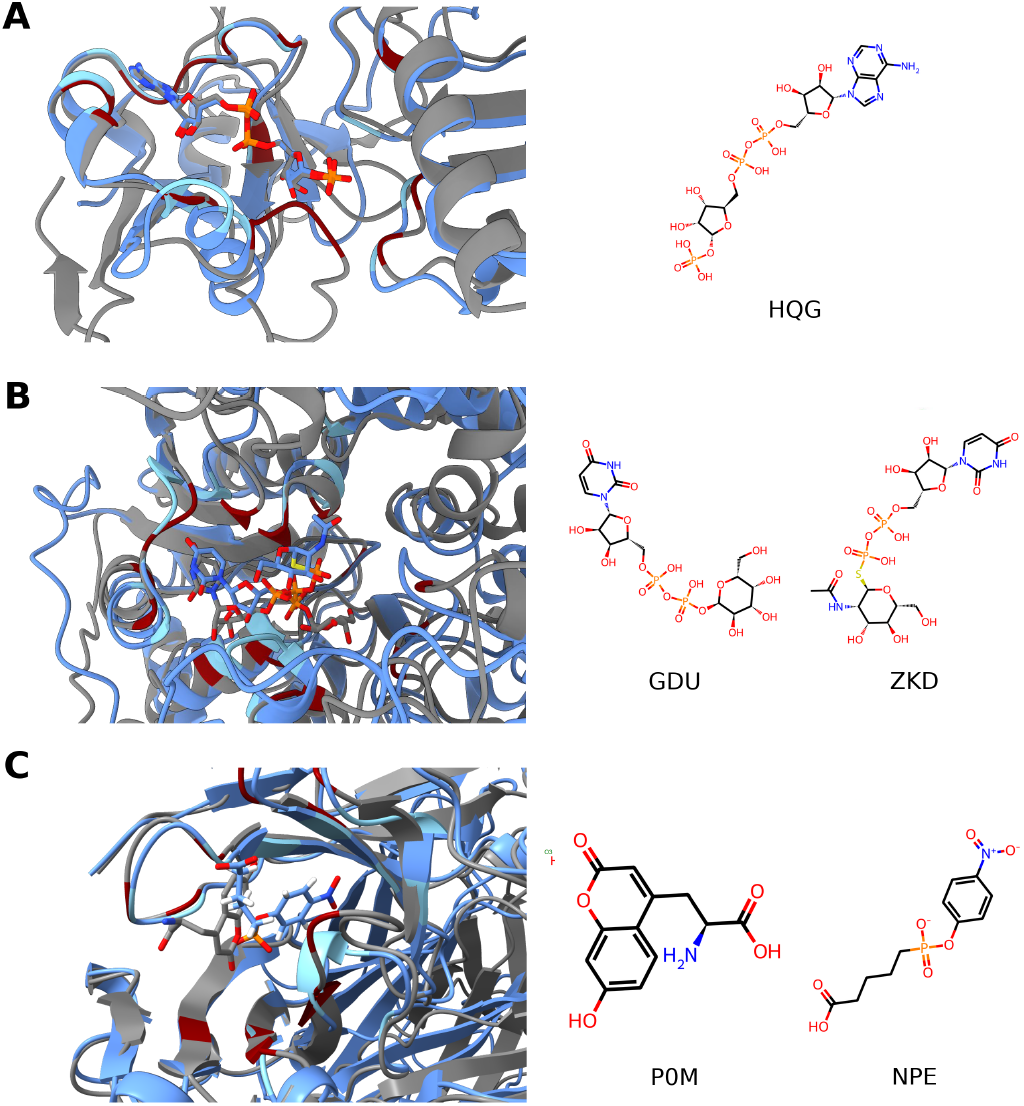
Binding structures of ‘False Negative’ cases. Ligand binding structures, binding ligands, and their binding sites for cases that are similar according to ProBiS but not similar according to LEN-Seek. Proteins in the evaluation data and their ligands are colored gray, while proteins in the latent space database are colored blue; amino acid residues of each ligand binding site are marked in red and sky blue, respectively. Of the two ligands on the right, the left one binds to the protein in the evaluation data, and the right one binds to the training data. A. Ligand binding structures for tRNA 2’-phosphotransferase (PDB ID: 7YW2, gray) and tRNA 2’-phosphotransferase (PDB ID: 6EDE, blue). Both bind the identical ligand HQG with similar patterns, but there are differences in detailed structure near the ligand binding site. B. Ligand binding structures for the glycosyltransferase WaaG from *Pseudomonas aeruginosa* (PDB ID: 8B62, gray; bound to UDP-galactose, GDU) and the O-GlcNAc transferase XcOGT from *Xanthomonas campestris* (PDB ID: 2XGO, blue; bound to UDP-thio-*N*-acetylglucosamine, ZKD). The two ligands are both UDP-sugars and highly similar, yet differences in the surrounding fold shift the residues that constitute the binding site. C. Ligand binding structures for an anti-hapten Fab antibody bound to L-hydroxycoumarylalanine (PDB ID: 8B50, gray; ligand P0M) and the 48G7 esterolytic catalytic antibody Fab (PDB ID: 1GAF, blue; ligand NPE). Both are immunoglobulin Fab fragments sharing the antibody fold but binding chemically distinct haptens.

Since the Chamfer distance sums the nearest-neighbor distances over all points in the two point clouds, LEN-Seek is sensitive to local mismatches and may overestimate the overall dissimilarity. When the bound ligands were identical, differences in the surrounding protein fold added residues to specific regions of the binding site. As seen in Figure 6A, two proteins binding ligand HQG shared the overall binding structure and similar residues, but differences in protein folding altered the region that constitutes the binding site.

For similar but non-identical ligands (Figure 6B), the shared substructures adopted similar binding geometries, but the differing parts bound differently, changing the amino acid composition and the binding region. Even for different ligands (Figure 6C), the same regions showed distinct binding structures, again diverging in residue composition.

Because structurally distinct residues are added, their latent vectors occupy different positions in latent space owing to the altered relative geometry.

Consequently, LEN-Seek’s false negatives appear to arise from defining binding sites as the residues within a fixed distance of the ligand, combined with the limitations of the Chamfer distance. This sensitivity stems from the current encoding, in which each residue is an individual vector, so the model is governed by the collective distances of individual residues rather than the global binding-site context.

In contrast, the ProBiS algorithm finds common subgraphs (maximum cliques) between two binding-site structures. By focusing on the intersection rather than the whole, ProBiS can report high similarity when the core structures match, even if some regions differ, making it robust to such cases.

### Structural Robustness

Analysis of ‘False Positive’ results—where LEN-Seek indicated similarity but ProBiS did not—revealed numerous cases where ligand binding structures were similar but residue positions in the binding site differed (Figure 7).

**Figure 7.**
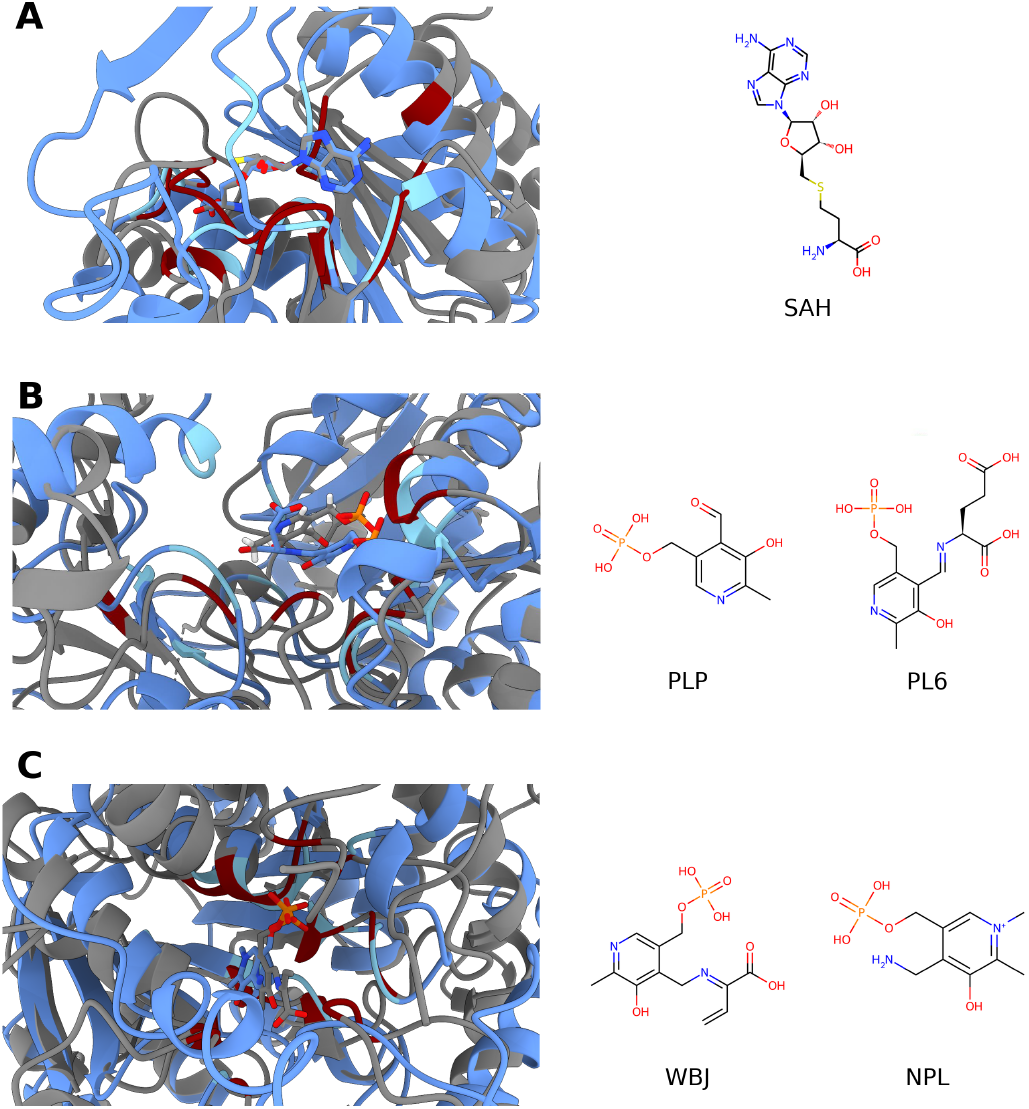
Binding structures of ‘False Positive’ cases. Ligand binding structures, binding ligands, and their binding sites for cases that are similar according to LEN-Seek but not similar according to ProBiS. While the binding structures are similar, the binding site itself is structurally ‘twisted’. A. Ligand binding structures for SpnK Methyltransferase (PDB ID: 8IAA, gray) and Mycobacterium hassiacum MeT1 (PDB ID: 6G80, blue). B. Ligand binding structures for F. varium tryptophanase (PDB ID: 8SIJ, gray) and E. coli Aspartate aminotransferase (PDB ID: 5VWR, blue). C. Ligand binding structures for Fub7 (PDB ID: 8ERB, gray) and E. coli Aspartate Aminotransferase (PDB ID: 1ASC, blue).

ProBiS considers alignment significant only if the RMSD of partial structure alignment is below a strict threshold (2.0 Å), so minute structural deformations can lead it to label a pair as dissimilar even when the two sites bind identical ligands. Because ProBiS is one reference method rather than absolute ground truth, we cross-checked these LEN-Seek-positive/ProBiS-negative pairs against an independent structural reference, G-LoSA. 77.9% (4,075 pairs) possessed a GA-score of 0.59 or higher (P-value *<* 0.05 [46]); that is, for most of these pairs the two reference methods disagree and an independent structural-alignment measure supports LEN-Seek’s similarity call rather than contradicting it.

### Impact of Feature Adaptation

Through the adaptation module, LEN-Seek transforms the 768-dimensional Ankh embedding space into an ‘adapted’ 512-dimensional space better suited to learning binding-site structure. This reduces the influence of the raw Ankh embeddings and strengthens the structural signal. To assess the impact of this adaptation, a ‘Non-adapted Model’ was trained to reconstruct the original 768-dimensional embeddings, thereby increasing the influence of the raw Ankh embeddings in latent space. Comparative analysis (Supplementary Figure S5, Supplementary Tables S4 and S5) showed that the non-adapted model had lower F1-score and MCC across all top-ranked ranges than the original model. Notably, the hit rate—the proportion of top-100 results that both LEN-Seek and ProBiS classify as similar—dropped sharply from 29.02% to 8.02%. The correlation with the G-LoSA GA-score also decreased (Supplementary Figure S6).

## Conclusion

We proposed LEN-Seek, a deep learning framework for high-speed retrieval of similar ligand binding sites in large-scale protein structure databases. By encoding the geometric and physicochemical context of a binding site—Ankh-derived residue features and roto-translational invariant edge geometry—into a probabilistic latent space with a Graph Transformer-driven VAE, it replaces costly pairwise alignment with distance computation in latent space, without equivariant networks or data augmentation.

Against ProBiS, LEN-Seek achieved a roughly 3,400-fold lower per-comparison cost (on its respective hardware) with high recall among top-ranked results. Its retrievals additionally agreed with ligand chemical similarity—the alignment-free criterion underlying template search—providing support from a measure independent of structural alignment. Its sensitivity to global residue composition, arising from residue-level encoding and the Chamfer distance, remains a limitation that future work could address by encoding each site into a single fixed-size latent vector via importance-aware pooling rather than naive averaging, by refining the latent space with metric learning[22], as in the margin-based contrastive objective of DeeplyTough[52], or by modeling binding-site flexibility through structural-uncertainty features such as B-factor[6] or AlphaFold PAE[27]. Overall, the learned latent space is a promising, scalable alternative for protein binding-site search and a useful building block for AI-driven drug discovery.

## Data availability

The source code for LEN-Seek is publicly available at https://github.com/arsenide33/LEN-Seek.

## Key Points

- Template-based binding-site search inherits the biological validity of experimentally determined sites, but pairwise structural alignment does not scale to databases of 10^5^–10^8^ structures.
- LEN-Seek encodes a binding site as a graph with protein language model node features and SE(3)-invariant edge geometry, so neither rotation augmentation nor an equivariant network is required.
- Retrieval reduces to a Chamfer distance between sets of latent vectors, giving a roughly 3,400-fold lower per-comparison cost than ProBiS.
- Retrievals agree with an alignment-free criterion: 88.0% of pairs called similar bind chemically similar ligands, versus 19.09% of pairs called dissimilar.
- Analysing where LEN-Seek and alignment-based tools disagree shows when a residue-level latent representation overestimates dissimilarity.

