## Supplementary Material for "LEN-Seek: Fast and scalable ligand binding-site similarity search in the latent space of an SE(3)-invariant graph VAE"

### Supplementary Figures

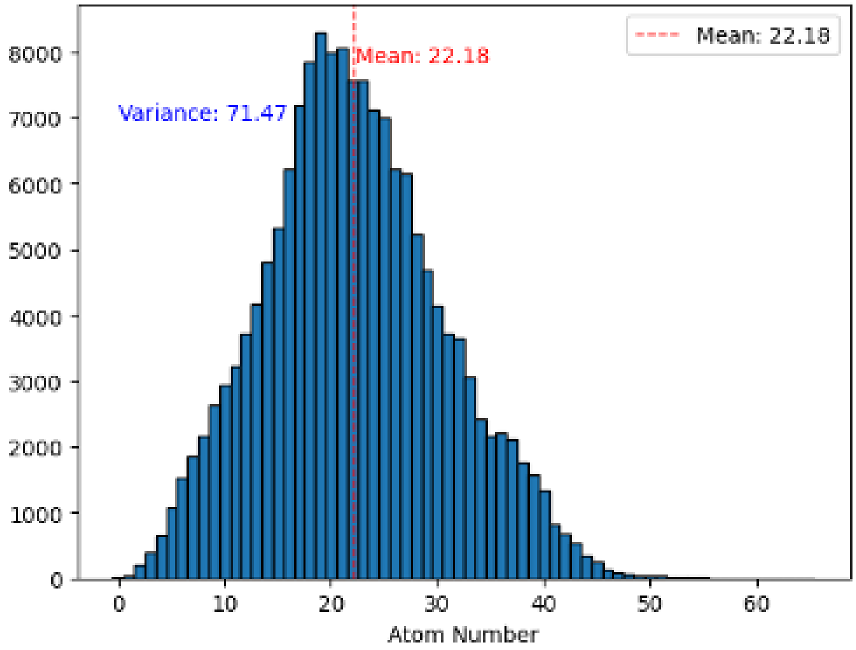


Figure S1: Statistics of the number of amino acids of each datapoint in BsitePDB. The maximum is 65 (mean 22.18, variance 71.47), which is the basis for unifying the number of graph nodes to 65.

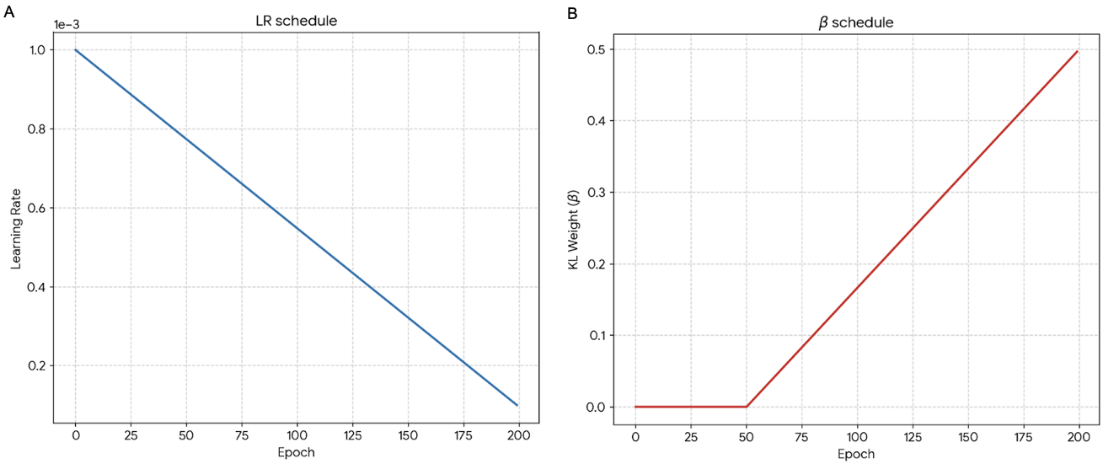


Figure S2: Epoch-wise change of learning rate and KL-divergence weight. (A) Learning rate starts at $10^{-3}$ and decays linearly to $10^{-4}$. (B) KL-divergence weight is set to 0 until epoch 50 for the model to focus on data reconstruction, then increases linearly.

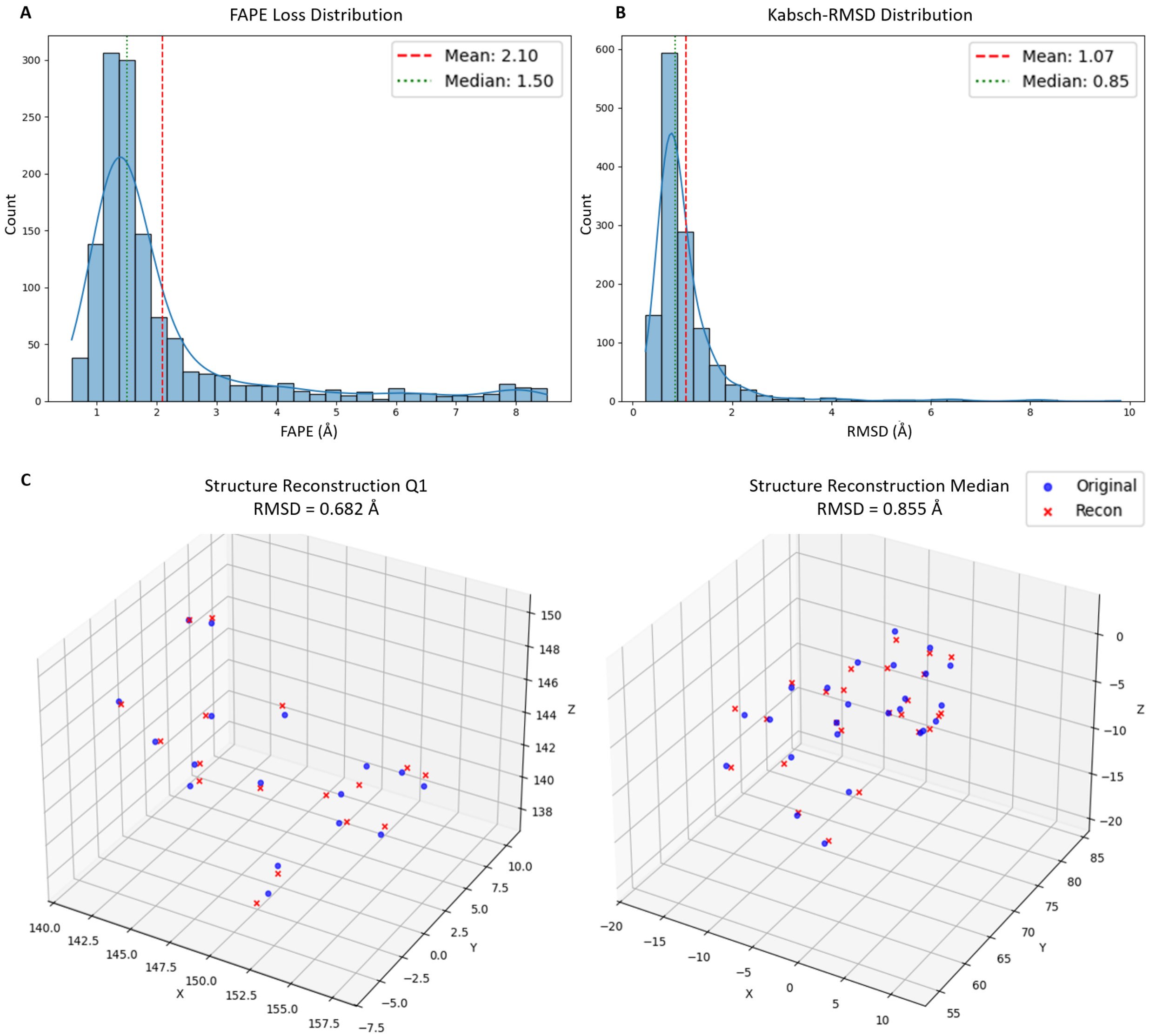


Figure S3: Structure reconstruction. (A) Distribution of FAPE between the original and reconstructed structures (mean 2.10 Å, median 1.50 Å). (B) Distribution of Kabsch RMSD between the original and reconstructed structures (mean 1.07 Å, median 0.85 Å). (C) Reconstructed versus original structures at the first-quartile (Q1) and median RMSD data points, with RMSDs of 0.682 and 0.855 Å, respectively. The original structure is shown in blue and the reconstructed structure in red.

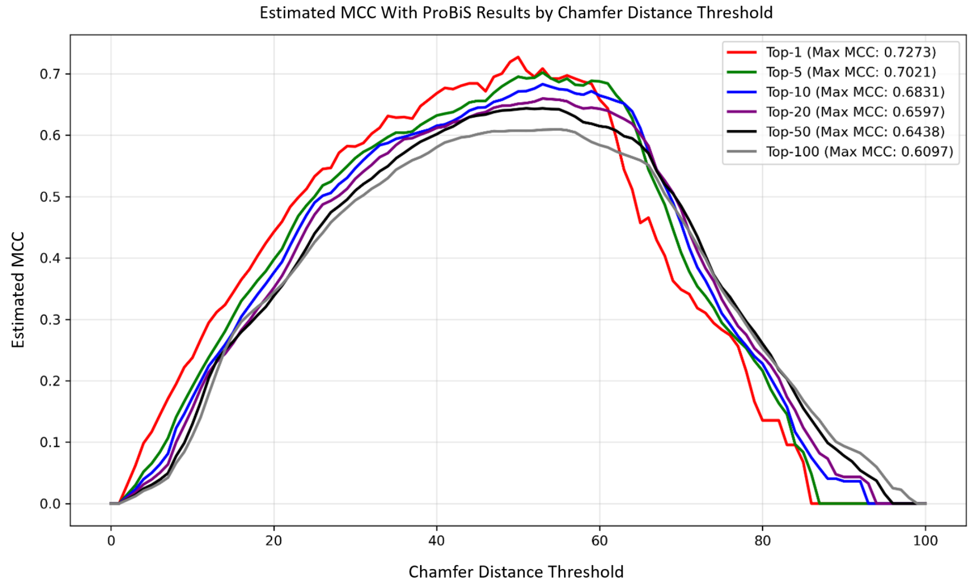


Figure S4: MCC variation by Chamfer-distance threshold for top-ranked results. Optimal values were 50 for top-1, 53 for top-5 to top-20, 51 for top-50, and 55 for top-100. A threshold of 50.0, yielding the maximum MCC for the top-1 result, was selected.

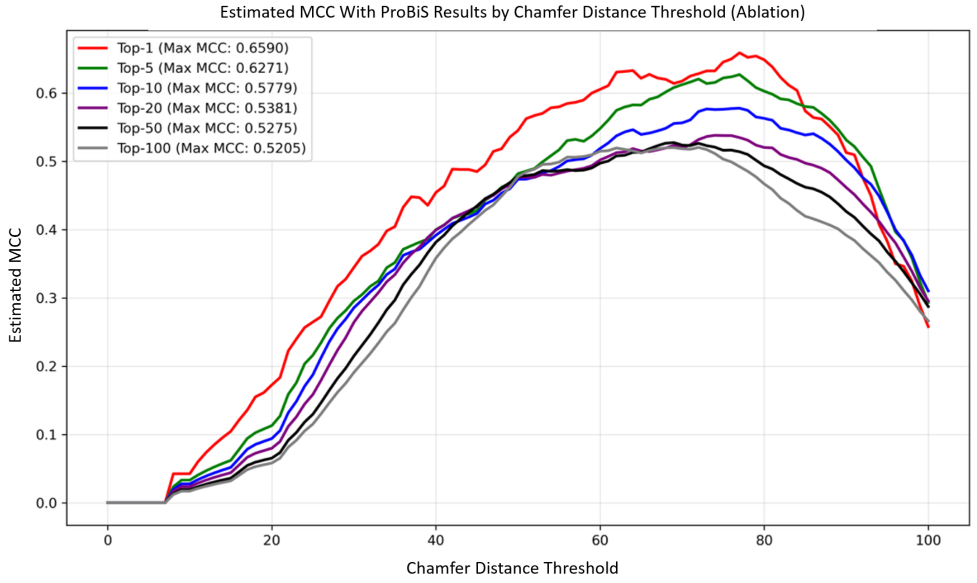


Figure S5: MCC variation by Chamfer-distance threshold for top-ranked results of the 'Non-adapted Model'. Optimal values were 77.0 for top-1, 74.0 for top-5 to top-20, 69.0 for top-50, and 72.0 for top-100. A threshold of 77.0, maximizing MCC for the top-1 result, was selected.

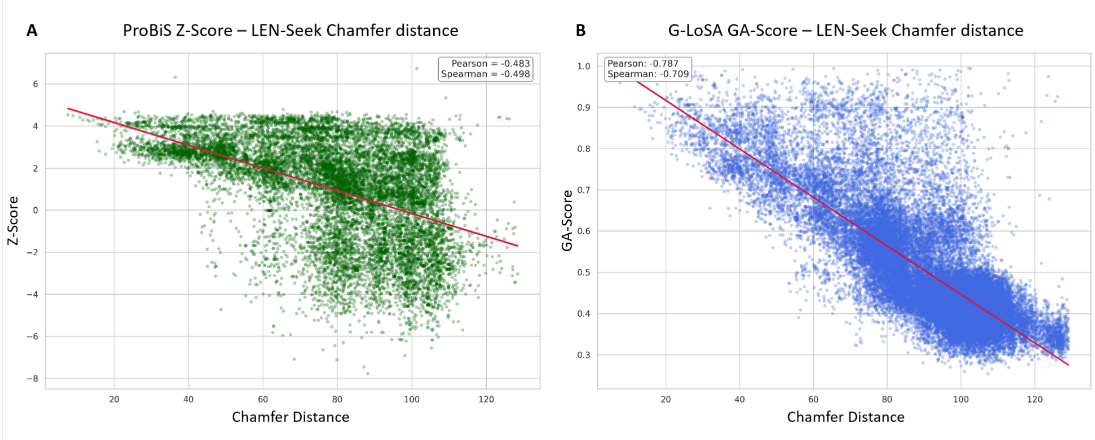


Figure S6: Score correlations of the top search results for the 'Non-adapted Model'. (A) Correlation between ProBiS Z-score and LEN-Seek Chamfer distance (Pearson −0.483, Spearman −0.498). (B) Correlation between G-LoSA GA-score and LEN-Seek Chamfer distance (Pearson −0.787, Spearman −0.709).

### Supplementary Tables

Table S1: Criteria of BsitePDB ligand binding site entry.

| No. | Criterion | Reference |
|---|---|---|
| 1 | Ligand must contain at least one carbon atom | sc-PDB |
| 2 | $140 \leq$ Ligand molecular weight $< 800$ | sc-PDB |
| 3 | No covalent bonds with protein | sc-PDB |
| 4 | Removal of water molecules not involved in $\geq 2$ hydrogen bonds in protein–ligand interaction | sc-PDB |
| 5 | Removal of molecules experimentally proven not to bind | ProBiS-Database |
| 6 | Distance from ligand heavy atoms to protein heavy atoms within (ligand heavy-atom van der Waals radius $+ 4$ Å) | ProBiS-Database |

Table S2: Hyperparameter setting.

| Hyperparameter | Meaning | Value |
|---|---|---|
| EPOCHS | Total training epoch | 200 |
| KL_START | Epoch to include KL divergence loss | 50 |
| START_LR | Learning rate of first epoch | 0.001 |
| END_LR | Learning rate of last epoch | 0.0001 |
| $\beta_{max}$ | KL divergence weight of last epoch | 0.5 |
| $\delta_{free}$ | KL divergence loss lower bound | 12 |
| $H$ | Number of heads in graph transformer | 8 |
| $L$ | Number of Graph Transformer layers | 4 |
| DROPOUT | Dropout rate | 0.1 |
| K_NEIGHBORS | Number of nodes to exchange information | 15 |
| $\lambda_{KL}$ | Weight of KL divergence loss | 1.0 |
| $\lambda_{MSE}$ | Weight of node feature value loss | 1.0 |
| $\lambda_{Cos}$ | Weight of node feature direction loss | 40.0 |
| $\lambda_{Norm}$ | Weight of node feature scale loss | 4.0 |
| $\lambda_{FAPE}$ | Weight of structure FAPE loss | 20.0 |
| $\lambda_{RMSD}$ | Weight of structure RMSD loss | 5.0 |

Table S3: Ratios by ligand similarity for LEN-Seek and ProBiS classification results. Analysis performed on the top-100 similarity results for each ligand binding site after screening the entire dataset with ProBiS and LEN-Seek, respectively. For ProBiS, a Z-score $\geq 2.0$ was set as 'Similar'; for LEN-Seek, a Chamfer distance $\leq 50.0$ was set as 'Similar'. Morgan-fingerprint Dice similarity of binding ligands was binned into 10 intervals.

| Ligand Dice Similarity | 0.0–0.1 | 0.1–0.2 | 0.2–0.3 | 0.3–0.4 | 0.4–0.5 | 0.5–0.6 | 0.6–0.7 | 0.7–0.8 | 0.8–0.9 | 0.9–1.0 |
|---|---|---|---|---|---|---|---|---|---|---|
| ProBiS Similar (%) | 1.76 | 12.56 | 9.41 | 4.21 | 5.46 | 3.51 | 5.61 | 15.86 | 3.87 | 37.75 |
| ProBiS Dissimilar (%) | 12.56 | 57.52 | 12.40 | 3.31 | 1.85 | 3.58 | 2.19 | 1.62 | 0.29 | 2.28 |
| LEN-Seek Similar (%) | 0.43 | 2.94 | 2.48 | 3.07 | 4.12 | 2.34 | 8.15 | 18.00 | 6.58 | 51.91 |
| LEN-Seek Dissimilar (%) | 11.17 | 50.83 | 13.13 | 4.62 | 1.99 | 3.77 | 3.20 | 3.79 | 1.94 | 5.56 |

Table S4: Classification based on ProBiS criteria for top search results of the 'Non-adapted Model'.

| Rank | TP | FP | FN | TN |
|---|---|---|---|---|
| 1 | 229 | 53 | 29 | 171 |
| 5 | 865 | 278 | 171 | 1095 |
| 10 | 1328 | 597 | 365 | 2520 |
| 20 | 1906 | 1160 | 722 | 5832 |
| 50 | 3809 | 2480 | 1387 | 17170 |
| 100 | 3861 | 4018 | 2091 | 38131 |

Table S5: Statistics based on ProBiS criteria for top search results of the 'Non-adapted Model'.

| Rank | Precision | Recall | F1-Score | MCC |
|---|---|---|---|---|
| 1 | 0.8121 | 0.8876 | 0.8481 | 0.6590 |
| 5 | 0.7568 | 0.8349 | 0.7939 | 0.6271 |
| 10 | 0.6899 | 0.7844 | 0.7341 | 0.5779 |
| 20 | 0.6217 | 0.7253 | 0.6695 | 0.5349 |
| 50 | 0.6057 | 0.7331 | 0.6633 | 0.5676 |
| 100 | 0.4900 | 0.6487 | 0.5583 | 0.4923 |
